# Note: Updating the metadata of four misidentified samples in the DrosRTEC dataset

**DOI:** 10.1101/2021.01.26.428249

**Authors:** Joaquin C. B. Nunez, Margot Paris, Heather Machado, Maria Bogaerts, Josefa Gonzalez, Thomas Flatt, Marta Coronado, Martin Kapun, Paul Schmidt, Dmitri Petrov, Alan Bergland

## Abstract

This note details the consortium’s rationale behind its decision to modify the metadata for putatively misidentified European samples in the DrosRTEC dataset. In brief, we use PCA on published datasets from North America and Europe to generate phylogeographic clusters reflective of worldwide *D. melanogaster* demography. We used this PCA to train a DAPC model in order the predict the group membership. Our results indicate that 4 out of 73 samples were misclassified and the metadata was updated accordingly. These samples are a spring-fall pair from Spain and Austria.

## INTRODUCTION

This note documents the metadata update for four putatively misidentified samples in the DrosRTEC dataset. These samples are listed in National Center for Biotechnology Information (NCBI) under accession numbers and were modified on NCBI on Oct 26,2020 (**Table 1**):

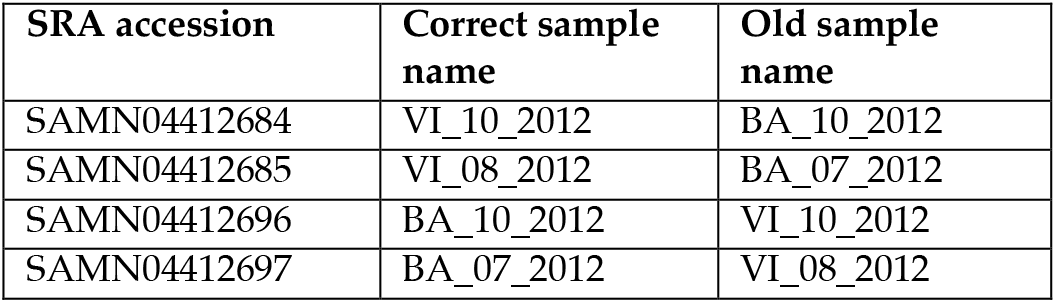

| SRA accession | Correct sample name | Old sample name |
| --- | --- | --- |
| SAMN04412684 | VI_10_2012 | BA_10_2012 |
| SAMN04412685 | VI_08_2012 | BA_07_2012 |
| SAMN04412696 | BA_10_2012 | VI_10_2012 |
| SAMN04412697 | BA_07_2012 | VI_08_2012 |

Inspection of hand-written notes shows that these samples were processed adjacent to each other during DNA extraction, and that a swap likely happened at this point. Here, we describe the procedures and rationale used in the NCBI record update.

## RESULTS & DISCUSSION

We used principal component analysis (PCA) to explore the potential ancestry backgrounds of the samples collected through the DrosRTEC consortium. Accordingly, we performed PCA using data from natural *Drosophila* populations collected from North America and Europe (Machado et al 2019; Kapun et al 2020). The PCA shows clear signatures of population structure in the first 3 PCs (PVE 18.4%, 5.1%, and 3.8%, respectively). We conducted a *k*-means clustering analysis on these 3 PCs to identify the major demographic groups of Europe and North America (**Fig 1: a-c**). We constrained the clustering algorithm to *k* = 3 groups based on the known demography for the species. This results in the identification of three phylogeographic clusters: North America (NoA; blue in **Fig 1**), Europe West (EUW; green in **Fig 1**), and Europe East (EUE; red in **Fig 1**). The split between Western and Eastern European fly populations has been noted before (Kapun et al 2020) in the DrosEU dataset. In that dataset, Spanish *D. melanogaster* populations cluster with the Western group (EUW), Ukrainian populations with the Eastern group (EUE), and Austrian fly populations are split between Western and Eastern cluster membership. Visual inspection of the principal components strongly suggests that the DrosRTEC European samples from Spain and Austria were mis-labeled: the DrosRTEC Spanish samples random SNPs to re-train the DAPC model. This analysis conservatively classifies all AT samples as part of the EUW cluster, and all ES samples as part clustered with the Eastern group, and the DrosRTEC Austrian with the Western group and reasonably close to other Spanish samples in PC space. The DrosRTEC Ukrainian samples appear to be properly labeled since they cluster in EUE PC space. These clustering patterns conspicuously suggest that samples from Spain (ES) and Austria (AT) were swapped.

**Figure 1:**
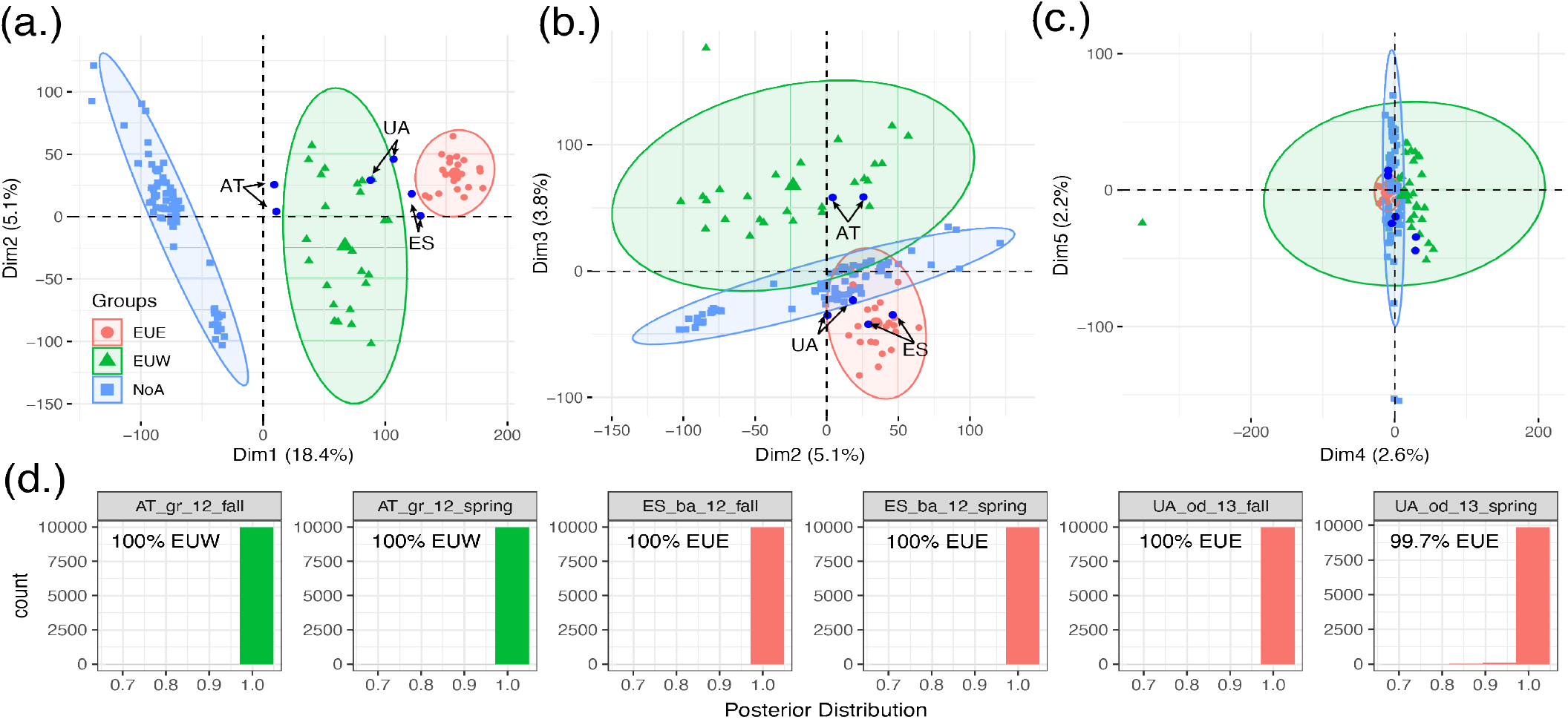
Evidence to substantiate the metadata update for misidentified samples. (a.) PCA components 1 and 2 of the joint DrosRTEC and DrosEU data. (b.) PCA components 2 and 3. (c.) PCA components 4 and 5. Across panels a-c: the blue samples indicate the misidentified samples added to the PCA as supplementary individuals. (d.) Posterior distribution of group membership for 10,000 iterations of the DAPC model. In addition to the posterior distribution, we annotated the name of the group membership of each sample, as well as the percentage of times the algorithm classified the sample into the group.

In order to assign group membership to each of the six misclassified samples, we trained a discriminant analysis of principal component (DAPC) model (Jombart, Devillard et al. 2010). We trained the DAPC model using the first 3 PCs from the PCA of the DrosRTEC and DrosEU data (**Fig 1: a-c**). We then used the DAPC model to estimate group membership posterior probabilities for each of the 6 European DrosRTEC samples. We repeated this process 10,000 times, each time sampling 10,000 of the EUE cluster. Lastly, all UA clusters are classified as part of the EUE cluster (**Fig 1d**).

## METHODS

All datasets used are publicly available: the DrosRTEC samples are discussed in Machado et al 2019. The DrosEU samples are discussed in Kapun et al. (2020). All analyses were done in the *R* (v. 4.0.0) statistical computing environment (R core team 2020). PCA was done using the *R* package FactoMiner v. 2.3 (Lê, Josse et al. 2008). Clustering analysis was done using the R package factoextra v. 1.0.7 (Kassambara and Mundt 2020). DAPC analysis was done with the R package adegenet v. 2.1.3 (Jombart and Ahmed 2011). Graphics were created using ggplot2 and other tools from the tidyverse (Wickham 2016, Wickham and Grolemund 2017).

